# Fluorescence and light scatter calibration allow comparisons of small particle data in standard units across different flow cytometry platforms and detector settings

**DOI:** 10.1101/796961

**Authors:** J.A. Welsh, J.C. Jones, V.A. Tang

## Abstract

Flow cytometers have been utilized for the analysis of submicron-sized particles since the late 1970s. Initially, virus analyses preceded extracellular vesicle (EV), which began in the 1990s. Despite decades of documented use, the lack of standardization in data reporting has resulted in a growing body of literature that cannot be easily interpreted, validated, or reproduced. This has made it difficult for objective assessments of both assays and instruments, in-turn leading to significant hindrances in scientific progress, specifically in the study of EVs, where the phenotypic analysis of these submicron-sized vesicles is becoming common-place in every biomedical field. Methods for fluorescence and light scatter standardization are well established and the reagents to perform these analyses are commercially available. However, fluorescence and light scatter calibration are not widely adopted by the small particle community as methods to standardize flow cytometry data. In this proof-of-concept study carried out as a resource for use at the CYTO2019 workshop, we demonstrate for the first-time simultaneous fluorescence and light scatter calibration of small particle data to show the ease and feasibility of this method for standardized flow cytometry data reporting. This data was acquired using standard configuration commercial flow cytometers, with commercially available materials, published methods, and freely available software tools. We show that application of light scatter, fluorescence, and concentration calibration can result in highly concordant data between flow cytometry platforms independent of instrument collection angle, gain/voltage settings, and flow rate; thus, providing a means of cross-comparison in standard units.

## Introduction

Efforts to standardize flow cytometry (FCM) data began several decades ago, following the production of multiple commercial flow cytometry platforms. Fluorescence standardization methods have been established since the late 1990’s, yet these have not been widely adopted by the FCM community for a variety of reasons. Nonetheless, extensive literature and commercially available materials have been developed to facilitate fluorescence calibration in cellular analysis^1–7^. These methods allow for the conversion of fluorescence intensities, which are arbitrary units, into standardized units of fluorescence recognized by the National Institutes of Standards and Technology (NIST), such as molecules of equivalent soluble fluorophore (MESF) or equivalent number of reference fluorophore (ERF)^1–7^. These protocols can be readily applied to the analysis of small particles, as demonstrated in this current study. Although it can be assumed that all commercial flow cytometers can fully resolve cellular populations from noise, the same is not true for small particles such as EVs. Therefore, supplementary methods, such as fluorescence calibration, need to be employed serve to validate small particle flow cytometry analysis.by quantifying the fluorescence signals such that data can be assessed empirically.

The detection and characterization of small particles, in the form of viruses, using light scatter triggering was published over 40 years ago^8^. Calibration of light scatter from a flow cytometer was demonstrated for small particles in 2009 by Fattacioli *et al*., and specifically for EVs by van der Pol *et al*. in 2012^9, 10^. Despite having been established for a decade, the use of light scatter calibration in small particle FCM has been limited, partly owing to the complexity of Mie Theory-based scatter modeling required for light scatter signal normalization. As an answer to this, in 2015, a commercial light scatter calibration assay (Rosetta Calibration by Exometry) was released in 2015 to facilitate this process and was used in a flow cytometry standardization study for the International Society on Thrombosis and Haemostasis (ISTH)^11^. In 2019, FCM_PASS_, a free alternative small particle flow cytometer calibration software package for light scatter and fluorescence became available^12^. While there is now both software and materials available for light scatter calibration, further support in the form of education and resource materials is required for the correct implementation and assessment in accuracy of calculated models.

Currently, both fluorescence and light scatter calibration is under-utilized in the field of small particle FCM. Calibration for small particle analysis is, however, critical due to the majority of commercial FCM instrumentation working at their detection limits when analyzing EVs and other biological particles <200 nm in diameter. Since flow cytometers have a wide range of optical configurations, methods are required for standardized data reporting such that meaningful biological conclusions can be made. There is currently no consensus method for this. This study was carried out for a CYTO2019 Workshop where the feasibility of combining scatter and fluorescence calibration for small particle FCM was presented^13^. Calibrations were performed using FCM_PASS_ software package with commercially available reference materials to convert fluorescence intensity to MESF and light scatter to diameter. A biological reference particle in the form of a fluorescently-tagged virus was used to validate this method^12^. Conversion of fluorescence and light scatter intensities from the antibody-labeled virus into PE MESF and nanometers allowed for direct comparison of the data from the same virus sample collected on two different flow cytometry platforms. This is the seminal report of the combined application of fluorescence and light scatter calibration as a method towards standardized data reporting for small particle FCM^12^.

## Materials and Methods

### Sample preparation

MV-M-sfGFP (ViroFlow Technologies, Canada, Lot#S1003A), murine leukemia virus tagged with super-folder GFP (sfGFP), was re-constituted and diluted as per manufacturer’s instructions according to the particle concentration provided. The virus particles were stained at a concentration of 5×10^8^ particles mL^−1^ and labeled with anti-GFP PE antibody (Clone FM264G, BioLegend) at 0.4 μg mL^−1^ in a 100 μL staining volume for a minimum of 30 minutes at room temperature, protected from light. To achieve this staining concentration, the lyophilized pellet of MV-M-sfGFP containing 1.8 x 10^9^ particles was re-suspended in the manufacturer recommended 200 μL volume of H_2_O and diluted to 10^9^ particles mL^−1^ by adding 1.6 mL of PBS. 50 μL of MV-M-sfGFP at 10^9^ particles mL^−1^ was then added to 50 μL of 0.8 μg mL^−1^ anti-GFP antibody (1 μL of 0.2 mg mL^−1^ antibody stock into 250 μL PBS). Upon completion of incubation, 2 μL of the labeled virus was diluted into 1 mL of PBS (~10^6^ particles mL^−1^) immediately prior to analysis by flow cytometry. Control samples such as antibody alone, MV-zero (virus expressing no GFP, Lot#Z1003A), and unstained MV-M-sfGFP samples were similarly prepared. All dilutions were made using 0.1 μm filtered PBS (PBS 1x, no Ca^2+^, no Mg^2+^, Wisent).

### Cytometer Configuration

BD LSR Fortessa 50 mW 488 nm laser, 50 mW 561 nm, 488/10 (SSC), 561-586/15 (PE), 488-530/30 (GFP). Beckman Coulter CytoFLEX S, 80 mW 405 nm, 50 mW 561 nm, 405/8 (SSC), 561-585/42 (PE), 488-525/40 (GFP). Further details on cytometer settings and acquisition can be found in attached MIFlowCyt (**Supplementary Table 1**) and MIFlowCyt-EV (**Supplementary Table 2**) documents^14, 15^.

### Small particle sample acquisition

The same bead and antibody labeled virus samples were acquired on an LSR Fortessa (BD Biosciences, USA) and CytoFLEX S (Beckman Coulter, USA). Samples were acquired using three different detection settings (for SSC and PE) to demonstrate that calibration can normalize data independently of acquisition settings and flow cytometry platform. The settings and acquisition plots are summarized in **Table 1** and **Supplementary Figure 1**, respectively. The detector settings for fluorescence detection on the CytoFLEX S and LSR Fortessa were determined using the PE MESF beads (QuantiBrite, BD Biosciences, USA). On the Fortessa, setting 1 being the lowest voltage where the dimmest bead population was resolved from background and setting 3 being the highest voltage where the brightest bead population was within maximum detection limit. Setting 2 was approximately halfway between setting 1 and 3. Gain settings for fluorescence detection on the CytoFLEX S were similarly determined. For scatter calibration three separate gains were used on the CytoFLEX S. For setting 1, side scatter (SSC) gain was chosen where the 600 nm NIST-traceable polystyrene bead population was within the range of detection. Setting 2 and setting 3 SSC voltages were the same for all three settings acquired on the LSR Fortessa. This was due to the LSR Fortessa being unable to fully resolve the virus above the trigger threshold. Since the concentration and comparison of fluorescent data would be heavily influenced by the SSC triggering threshold, a single voltage and trigger threshold was maintained. This provided a consistently detected population to compare across different fluorescent channel settings. The difference between the three scatter data sets on the CytoFLEX S are based on gating of MV-M-sfGFP at 20 and 50 PE MESF as outlined in the respective figures. All samples were acquired on the lowest preset instrument flow rates for one minute. For the CytoFLEX S this was 10 μL min^−1^, verified using the built in calibration software and weighing samples of deionized water before and after acquisition. For the LSR Fortessa the flow rate was calculated to be approximately 18.2 μL min^−1^ by measuring spike-in beads whose concentration was determined using the CytoFLEX S.

### Light scatter calibration

81, 100, 152, 203, 269, 303, 345, 401, 453, 568, 600 nm polystyrene NIST-traceable beads (ThermoFisher Scientific, USA) and 480 and 730 nm silica NIST-traceable beads (ThermoFisher Scientific, USA) were acquired on each cytometer. Median SSC-H intensity (488 nm SSC on LSR Fortessa, 405 nm SSC-H on CytoFLEX S) were gated using FlowJo (v10.5.3, USA). Mie modeling and subsequent conversion of light scatter intensity to diameter was performed using FCM_PASS_ software (v2.17, http://nanopass.ccr.cancer.gov)^12^. The background of how Mie theory and the FCM_PASS_ software work is beyond the scope of this technical note and can be found in previously published literature^16, 17^. Model input settings used including refractive indices, bead information, and statistics can be found in **Supplementary Table 3** and **4**, with model outputs shown in **Supplementary Figure 2** and **3**. The effective refractive index (RI) of the MV-M-sfGFP virus was assumed to be 1.45 based on published literature and dispersion was accounted for using Sellmeier equations for water^18, 19^. Samples were acquired with one scatter detector setting on the LSR Fortessa while 3 different scatter detector settings were acquired on the CytoFLEX S. A single bead population (152 nm) was used to cross-calibrate detector settings 2 with settings 1 and 3 on the CytoFLEX S with the full set of light scatter calibration beads acquired using detector setting 1.

### Fluorescence calibration

Linear regression was performed by converting acquired PE channel statistics and PE MESF reference bead values (QuantiBrite PE, Cat# 340495, Lot 73318, BD Biosciences, USA) to logarithmic values before performing regression, **Supplementary Figure 2D-F; Supplementary Figure 3D-F**. PE MESF reference beads were acquired at a single voltage/gain on each instrument that allowed for the brightest bead to be within the range of detection of the instrument. To account for differing spectral filters and as a demonstration of utilizing ergonomic calibration methods, 8-peak rainbow beads (Cat# RCP-30-5A, Lot AF01, Spherotech, USA) were cross-calibrated to PE MESF values on each cytometer at the same voltage/gain as the PE MESF beads were acquired at. The derived MESF values are summarized in **Supplementary Table 5.** Median PE signals for each population were gated using FlowJo. The linear regressions for fluorescence calibration and conversion of .fcs files to MESF units was performed using FCM_PASS_ software (v2.17, http://nanopass.ccr.cancer.gov)^12^. The cross-calibrated 8-peak rainbow beads were used to calibrate the PE intensity scales at detector settings 2 and 3 for both the LSR Fortessa and settings 1 and 3 for the CytoFLEX S.

### Concentration calibration

Virus concentration on the CytoFLEX S and LSR Fortessa was normalized using 200 nm fluorescent polystyrene (Green FluoSpheres, Cat# F8848, ThermoFisher Scientific, USA) spike-in beads, which were mixed with the virus sample to evaluate relative bead to virus counts. The mean concentration of beads was determined using the CytoFLEX S post-calibration of the fluidic system at a low flow rate. Due to composition of the spike-in beads, aggregates were observed. Populations appearing as doublets, triplets, etc, were counted as a single bead.

### Data and Statistical Analysis

All data analyses and figures were carried out using MATLAB (R2019b, Mathworks, Inc) unless otherwise stated. Samples were gated as shown in representative example **Figure 1** and exhaustively in **Supplementary Figure 1** with control samples shown in **Supplementary Figure 4**.

**Figure 1.**
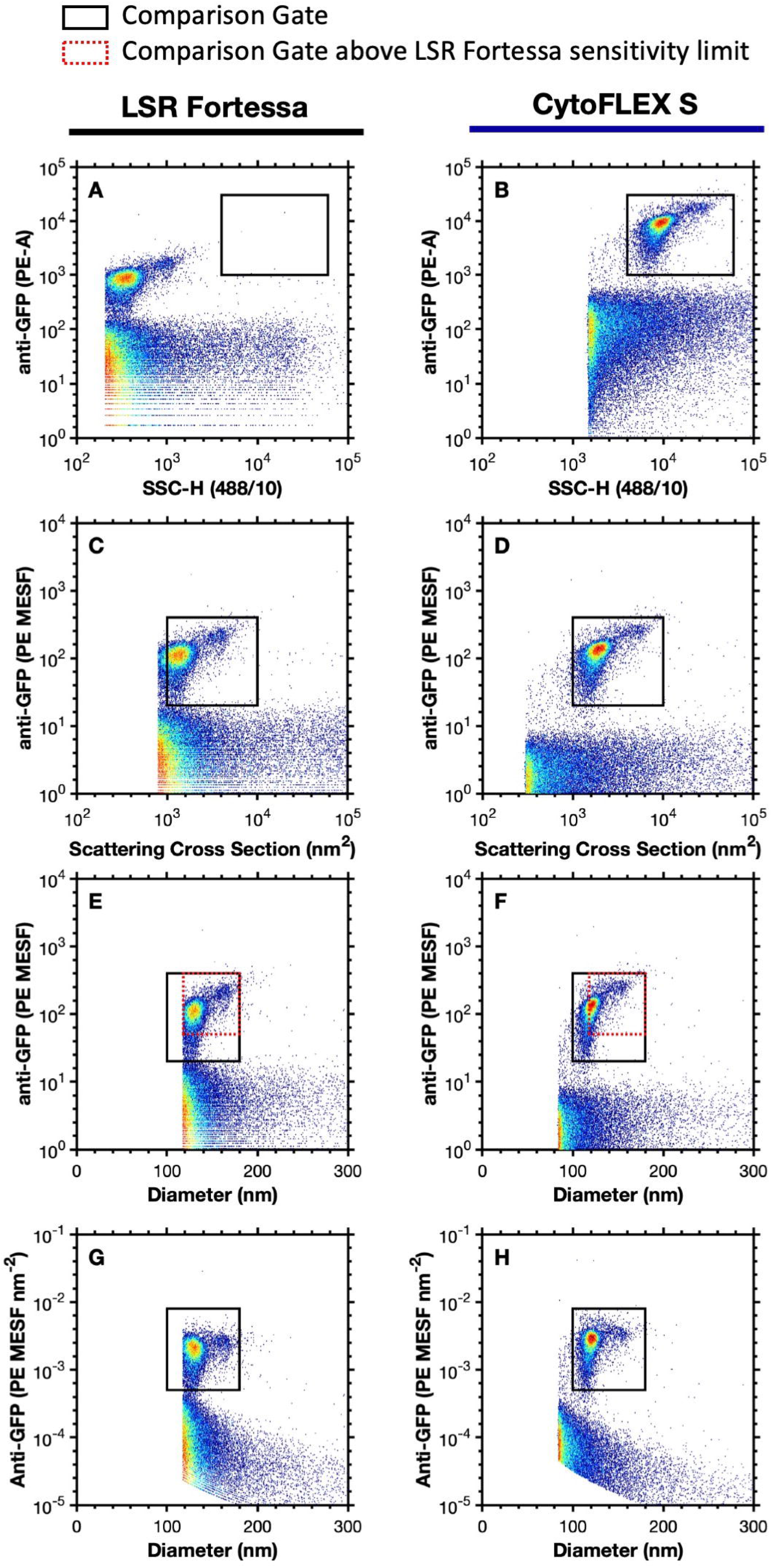
Representative data demonstrating the comparability of uncalibrated and calibrated data across flow cytometers. **A-B)** The side scatter versus anti-GFP-PE fluorescence of MV-M-sfGFP are shown on the LSR Fortessa and CytoFLEX S, respectively. A comparison gate is drawn from 4 x 10^3^ to 6 x 10^4^ SSC-H and 1 x 10^3^ to 3 x 10^4^ PE-A. **C-D)** The scatter cross section versus Anti-GFP-PE (MESF) of MV-M-sfGFP are shown on the LSR Fortessa and CytoFLEX S, respectively. A comparison gate is drawn from 1 x 10^3^ to 1 x 10^4^ nm^2^ and 20 to 400 PE MESF. **E-F)** The diameter (nm) versus Anti-GFP-PE (MESF) of MV-M-sfGFP are shown on the LSR Fortessa and CytoFLEX S, respectively. A comparison gate is drawn from 100 to 180 nm to and 20 to 400 PE MESF (black). A second comparison gate is drawn from 118 to 180 nm to and 50 to 400 PE MESF (red dotted). **G-H)** The diameter (nm) versus Anti-GFP-PE (MESF nm^−2^) of MV-M-sfGFP are shown on the LSR Fortessa and CytoFLEX S, respectively. A comparison gate is drawn from 100 to 180 nm to and 5 x 10^−4^ to 8 x 10^−3^ PE MESF nm^−2^ (black).

## Results

### Fluorescence calibration

The median PE intensity of anti-GFP-PE labeled MV-M-sfGFP from the LSR Fortessa and CytoFLEX S (gated from 50-400 PE MESF) ranged in arbitrary units of intensity from 111 to 17604 (a 159-fold difference), **Table 2**, across the three different detectors settings per cytometer, **Table 1**. The result of this was that a single gate could not be used irrespective of instruments or detector settings, **Figure 1A-B, Figure 2B, Supplementary Figure 1A-F**. Upon calibration of fluorescence intensity to PE MESF units, median values ranged from 99 to 128 PE MESF (29% difference) across flow cytometer platforms. Within cytometry platforms this variation was seen to be 103 to 110 (7% difference) and 99 to 128 (29% difference) PE MESF for the LSR Fortessa and CytoFLEX S, respectively, **Table 2**. Variation within and between flow cytometry platforms irrespective of detector settings was therefore considerably more consistent upon fluorescence calibration. Furthermore, a single gate from 20 to 400 PE MESF could be used to gate labelled MV-M-sfGFP irrespective of instrument or detector settings, **Figure 2E**.

**Figure 2.**
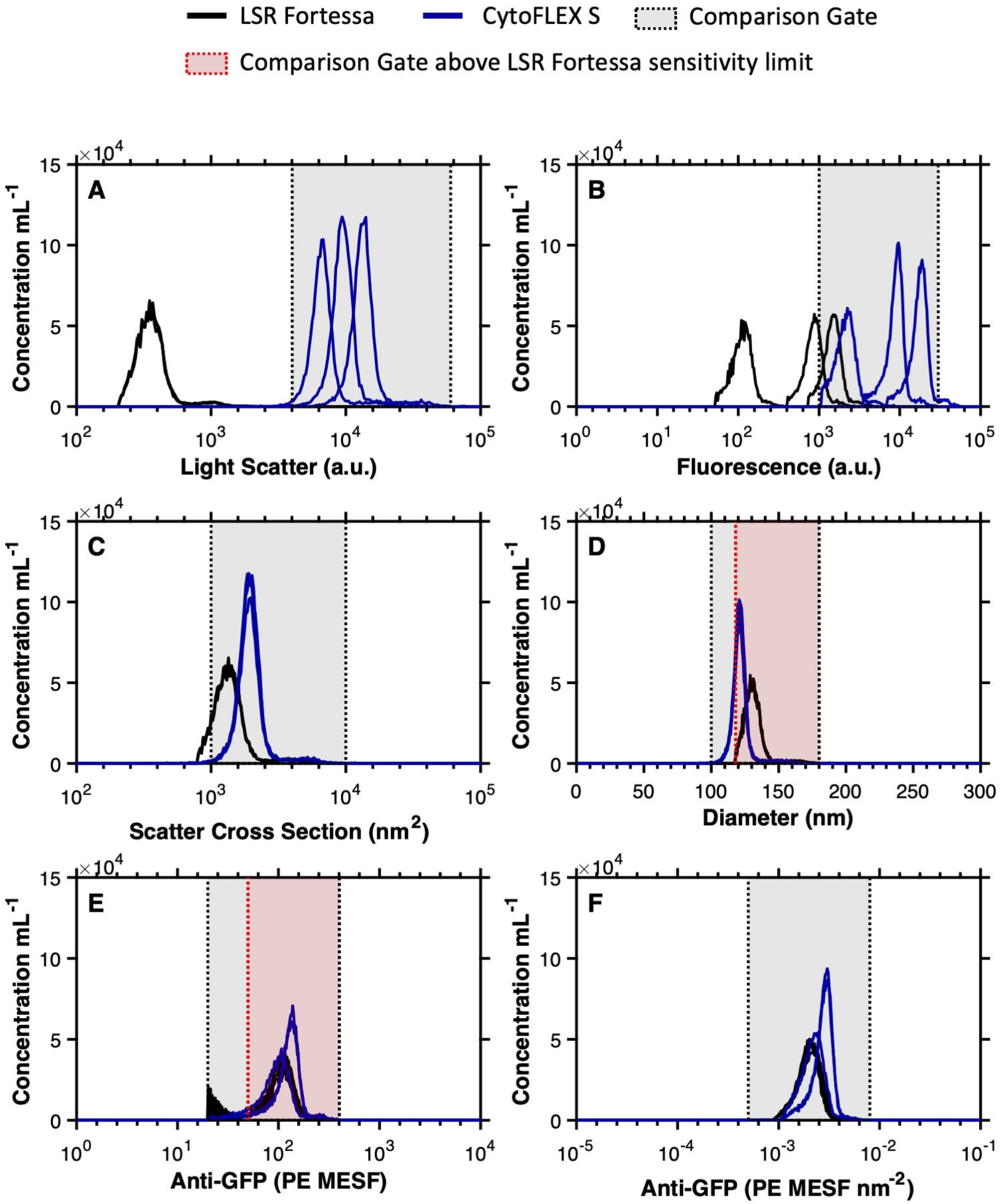
Overlays of uncalibrated vs. calibrated data from anti-GFP-PE labeled MV-M-sfGFP on flow cytometers at different detector settings. **A)** Uncalibrated light scatter intensity of anti-GFP-PE labelled MV-M-sfGFP is shown on the LSR Fortessa (black) and CytoFLEX S (blue). A comparison gate is drawn from 4 x 10^3^ to 6 x 10^4^ SSC-H. **B)** Uncalibrated fluorescence intensity of anti-GFP-PE labelled MV-M-sfGFP is shown on the LSR Fortessa (black) and CytoFLEX S (blue). A comparison gate is drawn from 1 x 10^3^ to 3 x 10^4^ PE-A. **C)** Calibrated light scatter intensity in scatter cross section units of anti-GFP-PE labelled MV-M-sfGFP is shown on the LSR Fortessa (black) and CytoFLEX S (blue). A comparison gate is drawn from 1 x 10^3^ to 1 x 10^4^ nm^2^. **D)** Calibrated light scatter intensity in diameter units of anti-GFP-PE labelled MV-M-sfGFP is shown on the LSR Fortessa (black) and CytoFLEX S (blue). A comparison gate is drawn from 100 to 180 nm (grey) and a second comparison gate from 118 to 180 nm (red) is drawn to allow for comparisons above the LSR Fortessa SSC-H trigger threshold. **E)** Calibrated fluorescence intensity in PE MESF units of anti-GFP-PE labelled MV-M-sfGFP is shown on the LSR Fortessa (black) and CytoFLEX S (blue). A comparison gate is drawn from 20 to 400 PE MESF (grey) and a second comparison gate from 50 to 400 nm (red) is drawn to allow for comparisons above the background signal from the LSR Fortessa at low voltages. **F)** Calibrated fluorescence intensity in PE MESF nm^−2^ units of anti-GFP-PE labelled MV-M-sfGFP is shown on the LSR Fortessa (black) and CytoFLEX S (blue). A comparison gate is drawn from 5 x 10^−4^ to 8 x 10^−3^ PE MESF nm^−2^ (grey).

### Light scatter calibration

Side-scatter intensity of MV-M-sfGFP was compared between the LSR Fortessa and CytoFLEX S. Three scatter settings were tested on the CytoFLEX S and one setting on the LSR Fortessa (defined in methods section and **Table 1**). The SSC signal of the MV-M-sfGFP virus between instruments ranged from 347 to 13121 arbitrary units (a 37.8-fold difference) prior to calibration, **Table 2**. The result of this was that a single gate could not be used irrespective of instruments or detector settings, **Figure 1A-B, Figure 2A, Supplementary Figure 1A-F**.

While scattering cross section is a valid metric to report calibrated light scatter measurements, it describes the statistical quantity of light that will reach the detector. This metric is influenced by the collection optics of the instrument, namely that larger collection angles will collect more light. The light scatter cross section can, therefore, normalize results from instruments with the same collection angle irrespective of detector settings, as seen in **Figure 2C.** Upon calibration of CytoFLEX S SSC signal from arbitrary units to scatter cross section the range reduced from a 37.8-fold difference to <1%. Scatter cross section as a unit cannot, however, normalize data between instruments with different collection angles. If the same population is acquired on two instruments with differing collection angles, the instrument with the larger collection angle will most likely result in a larger scattering cross section. This is observed when the CytoFLEX S and LSR Fortessa data are compared, **Figure 1C-D, Figure 2C**. The scatter cross-section of the MV-M-sfGFP virus on the CytoFLEX S (collection half angle of 53.2°) was 1903 nm^2^, while the LSR Fortessa (collection half-angle of 46.6°) had a scattering cross-section of 1323 nm^2^ (a difference of 44%) **Table 2**.

To normalized light scatter signals irrespective of collection angle a scatter-diameter curve must be generated (**Supplementary Figure 2C, 3C**) which takes into account the collection angle of the system. In order for the approximated diameter to be accurate, knowledge and incorporation of a suitable refractive index (RI) of the particles being normalized is required. It can be seen that when the MV-M-sfGFP data is converted to units of diameter, the range between the instruments is reduced to 121 nm - 130 nm (a 10.7% difference), **Table 2, Figure 1E-F, Figure 2D**. Within the CytoFLEX S platform the variation in derived diameter irrespective of detector settings was seen to be <0.2% with derived diameter varying from 120.6-120.8 nm, **Table 2**. The comparability of the data between uncalibrated and calibrated values across the two CytoFLEX S and LSR Fortessa can be seen in representative examples, **Figure 1, 2**, and for every setting (**Supplementary Figure 1**). Furthermore, a single gate from 100-180 nm can be used irrespective of instrument and detector settings to identify the virus population, **Figure 2D**.

### Epitope-density/MESF-density calibration

Epitope-density describes the number of the epitopes over a unit of surface area e.g. epitope number nm^−2^. An alternative, in cases where there is more than one fluorophore per antibody, could be the number of fluorophores over unit of surface area e.g. MESF nm^−2^. The epitope number can be considered equivalent to the MESF values obtained from fluorescence calibration when the following criteria are met: 1) fluorophore to protein (F:P) ratio is 1:1^20^. Due to the large size of PE, this can generally be assumed when conjugated IgG, whereas small dyes such as fluorescein can vary considerably. 2) All antibodies have the same dye. Tandem dyes for example can result in variability of fluorescence due to the tandem not being present on all antibodies. 3) Antibody-labeling is optimized i.e. all epitopes are labelled. 4) One antibody is bound per one epitope, and 5) no steric hindrance impeding the antibody from binding to all expressed epitopes.

When compared to MV-M-sfGFP, EVs have a very wide diameter distribution (**Figure 4A**) which is proportional to their surface area distribution (**Figure 4B**). If the surface area of EVs scale with their epitope number, this too will be highly variable (**Figure 4C-D**)^21–23^. EVs have a lower RI and the majority are likely smaller than MV-M-sfGFP, resulting in decreased light scatter detectability. While high abundance epitopes can exist on EVs, many epitopes are likely much lower than GFP-expression on MV-M-sfGFP^22, 23^. Statistics, such as a median or mean, for reporting fluorescence and diameter are heavily dependent on how much of a population is being resolved. Therefore, if a population is not fully resolved the reporting of any statistic to describe the population in terms of light scatter, fluorescence, or concentration parameters requires it to accompanied by the limit of detection to be reproducible e.g. 3 x 10^6^ particles mL^−1^ (30-300 PE MESF).

Upon calibration of fluorescence intensity to PE MESF nm^−2^ units, median values ranged from 2.02 x 10^−3^ to 2.78 x 10^−3^ PE MESF nm^−2^ (38% difference) across flow cytometer platforms. Within cytometry platforms this variation was seen to be 1.93 x 10^−3^ to 2.06 x 10^−3^ (7% difference) and 2.16 x 10^−3^ to 2.80 x 10^−3^ (30% difference) PE MESF for the LSR Fortessa and CytoFLEX S, respectively, **Table 2**. Variation within and between flow cytometry platforms irrespective of detector settings was therefore considerably more consistent using PE MESF nm^−2^ than no calibration, with variation similar to that of MESF calibration, **Figure 1G-H, Figure 2F**.

The epitope- or MESF-density may potentially provide a useful metric to allow better differentiation and consistent reporting of epitope expression. This statistic removes variability that arises due to heterogeneity of surface area in samples where there is a large size range (e.g. EVs) and available instrumentation will have differing limits of detection. Given the current challenges to fully resolve all EVs in a sample, epitope density offers a means to report on expression level per particle when knowledge of expression within the whole population is not yet feasible. To further illustrate this point, a hypothetical example of surface marker expression on EVs is shown with values reported in epitope number in comparison with epitope density (**Figure 4D-E**), assuming the limit of detection is 20 PE MESF (**Figure 4D**, red dashed line). There can be significant overlap in expression between the example markers when simply reported as epitope number, while minimal overlap is seen in epitope density populations despite having a coefficient of variation of ±20%. Provided the epitope density scales consistently with the population, it is therefore feasible that the epitope/MESF-density would be consistent even if only a portion of the population is detectable. This would allow a much more consistent method of reporting and comparing data between instruments with differing limits of detection, although further validation would be required. This metric has not previously been demonstrated for small particles and does require care in implementation and interpretation due to potential inaccuracies. The reliability of epitope/MESF-density as a statistic will be dependent upon the precision of the MESF and diameter derivation methods.

### Concentration calibration

The technical specifications for the low flow rate on the LSR Fortessa is 12 μL min^−1^, while the lowest calibrated flow rate on the CytoFLEX S is 10 μL min^−1^. Using spike-in beads, the mean flow rate was found to be 18.2 μL min^−1^ on the LSR Fortessa. The recorded number of MV-M-sfGFP events if fully detected should theoretically have been 20% higher than those of the CytoFLEX S. In practice, the mean total count difference between instruments was 39%, **Table 2, Figure 3A**. This indicates that either the flow rate between the two instruments is not as stated, or the virus was resolved to different extents between the instruments, or a combination of the two. Differences between advertised and observed flow rates has previously been reported in small particle standardization studies^11^.

**Figure 3.**
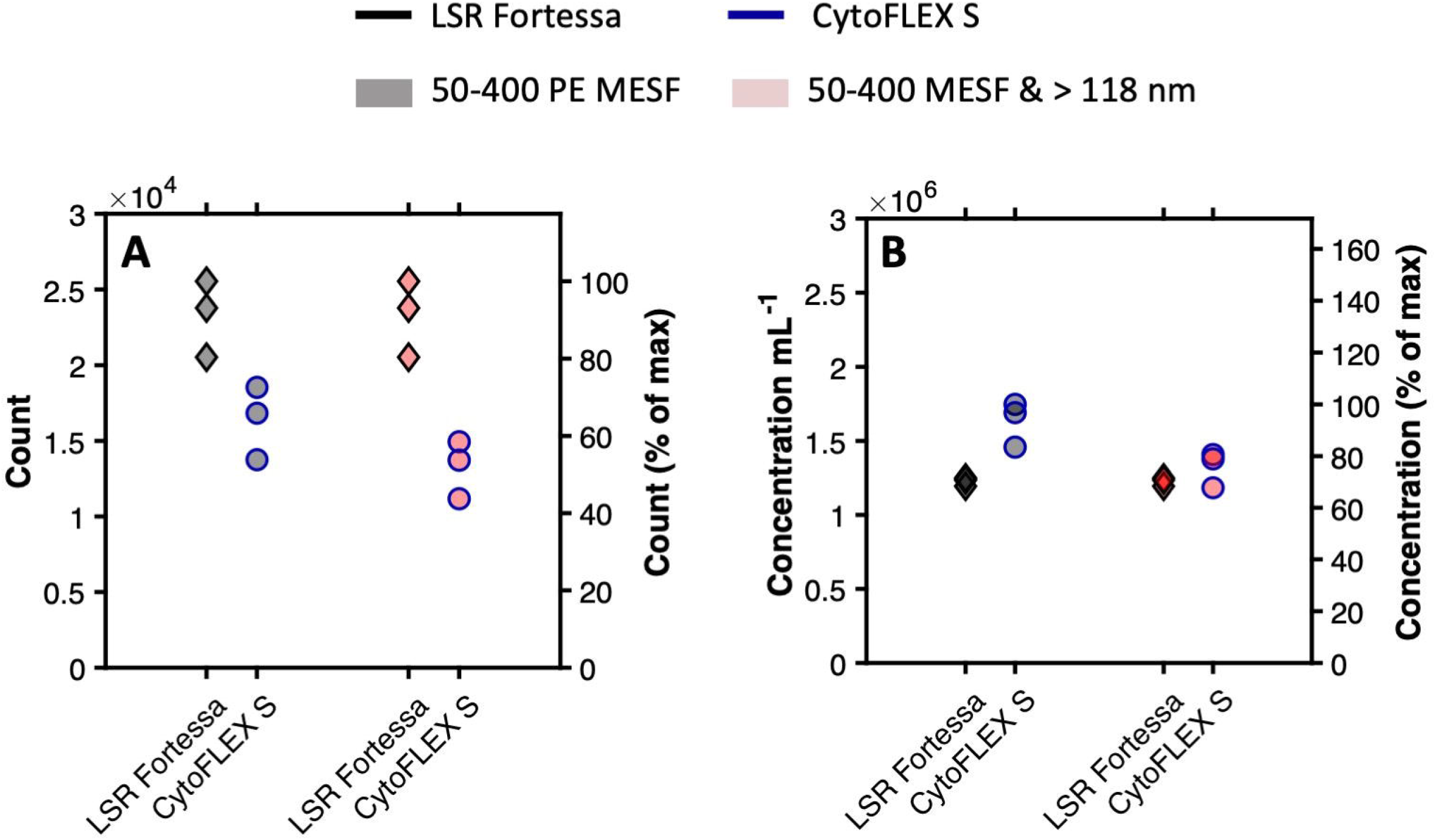
Comparisons of uncalibrated vs. calibrated abundance of detected anti-GFP-PE labeled MV-M-sfGFP on flow cytometers at different detector settings. **A)** comparison of acquired counts in 1 minute on LSR Fortessa (black outline) and CytoFLEX S (blue outline) gated from 50-400 PE MESF units (grey symbol) and 50-400 PE MESF and >118 nm (red symbol). **B)** comparison of acquired concentration per mL in 1 minute on LSR Fortessa (black outline) and CytoFLEX S (blue outline) gated from 50-400 PE MESF units (grey symbol) and 50-400 PE MESF and >118 nm (red symbol).

**Figure 4 –.**
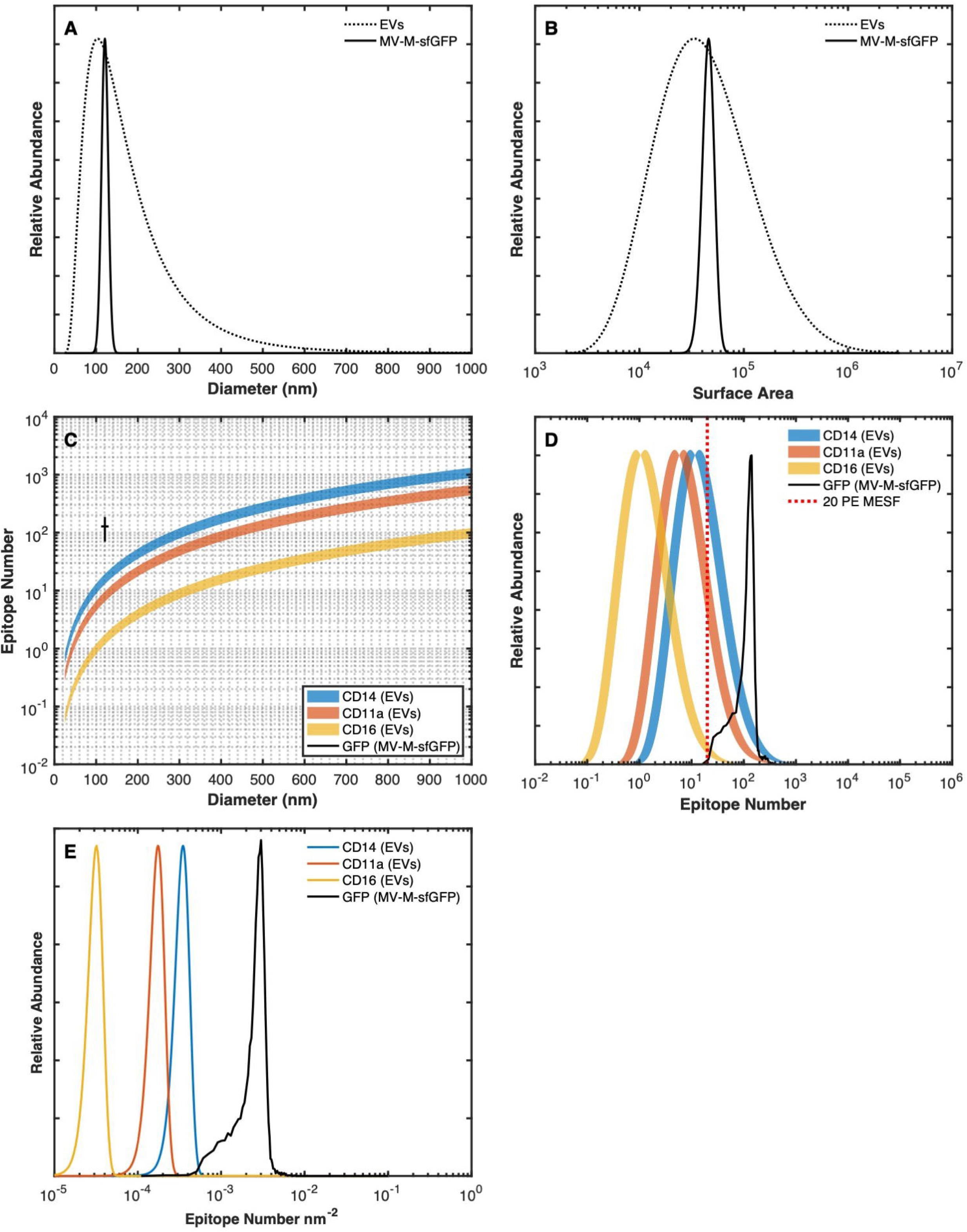
Reporting epitope abundance. **A)** The diameter distribution of extracellular vesicles based on the current literature (dotted), assuming a log-normal distribution with a modal point at ~100 nm, versus the diameter distribution of murine leukemia virus, 124±14 (solid)^21^. **B)** A comparison of surface area distribution of extracellular vesicles (dotted) versus murine leukemia virus based on the diameter distribution. **C)** Comparison of epitope number across EVs based on scaling previously published leukocyte (10 μm) epitope abundance for CD14 (110,000 epitopes) (blue), CD11a (55,000) (orange), and CD16 (10,000) (yellow)^22^. Each has a variation of ±20%. The calculated GFP number of MV-M-sfGFP assuming one phycoerythrin molecule per antibody, one antibody per GFP, no steric hindrance, and enumeration using PE MESF regression. MV-M-sfGFP epitope data was based on acquired data from the CytoFLEX S at Setting 2 (**Table 1**) is shown at 124±14 nm and 128±50 GFP epitopes. **D)** Demonstrates epitope number distribution based on the surface area data (**B**) and the epitope number vs. diameter plot (**C)**. **E)** Demonstrates reporting epitope-density distribution of each marker (CD14, CD11a, CD16, GFP) based on the data from plot **C**.

When spike-in beads were used to normalize the MV-M-sfGFP concentration between the two instruments, the calculated mean concentration was shown to be 33% higher on the CytoFLEX S (1.63 x 10^6^ particles mL^−1^) than on the LSR Fortessa (1.23 x 10^6^ particles mL^−1^), **Table 2, Figure 3B**. The inability of the LSR Fortessa to fully resolve MV-M-sfGFP is a possible source of the concentration discrepancy between the two cytometers. This can be observed on the SSC parameter’s triggering threshold channel at 200 on **Figure 1A, 2D** & **Supplementary Figure 1A-C**. To investigate if this accounts for the difference observed in concentration, a gate above the scatter trigger threshold of the LSR Fortessa was constructed from 50-400 PE MESF and 118-180 nm, **Figure 1E-F, 2D-E**. Using this gate, the difference in mean MV-M-sfGFP concentration on the

CytoFLEX S was reduced to just 7.8% at 1.32 x 10^6^ particle mL^−1^ versus 1.23 x 10^6^ particles mL^−1^, **Table 2, Figure 2E-F, 3A-B**.

While the spike-in beads used here demonstrate that concentration can provide effective normalization between platforms when flow cytometer sensitivities are accounted for through calibration, the accuracy of the concentration spike-in beads can vary. It was noted in this work that the choice of spike-in beads in the form of 200 nm FluoSpheres was sub-optimal due to their tendency to aggregate. Future experiments should utilize more stable spike-in beads for added confidence in concentration enumeration.

### Calibration overview

When uncalibrated MV-M-sfGFP fluorescence and light scattering data is overlaid between the CytoFLEX S and LSR Fortessa, very few comparisons can be made despite this being the same sample acquired on both instruments, **Figure 2A-B**. This is illustrative of the current state of much of the small particle literature. The use of any one of the calibration metrics (MESF units, scatter cross section, diameter, MESF nm^−2^) all considerably increased the comparability of data in contrast to uncalibrated data, **Figure 2C-F**. Not only has this work exhibited the utility of fluorescence and light scatter calibration, but the relative ease of these methods have also been demonstrated. Once an instrument has been initially calibrated using MESF reference materials and a set of light scatter reference materials, future instrument measurements and samples can be cross-calibrated using just two calibration samples; one for fluorescence (cross-calibrated 8-peak beads) and one for light scatter (a single light scatter reference bead). On stable instrument optical models this data can be transposed using a single standard due to the collection angle being unlikely to change. Once cross-calibrated on a single platform, 8-peak rainbow beads should also be sufficient for MESF calibration, assuming collection filter sets are not altered. While calibration can be very ergonomic as outlined and demonstrated by cross-calibrating, any time an instrument is modified or altered e.g. from a regular service, MESF and light scatter calibration should be conducted in full again. Furthermore, it is recommended that this calibration be done periodically for good practice and also to monitor instrument performance. Users may not always be aware of modifications made to core instruments and these calibrations would ideally have help from shared resource laboratory managers.

## Discussion

Two main factors contributed to the consistency of the results in this report. 1) The same validation sample (PE-labeled MV-M-sfGFP) was used for all of the data acquisitions. This eliminated inconsistencies in staining intensity and highlights the importance of accurate reporting of both antibody concentrations as well as particle concentrations used for antibody labeling experiments in small particle FCM. Discrepancies in either of these parameters greater than a factor of two can significantly impact staining intensity of small particles^24^. 2) The specific types of beads used to perform the calibrations were pivotal for this standardization process. For the light scatter calibration beads, it is critical that beads are accurately sized and have, preferably measured, refractive indices. Light scatter calibration is best done with non-fluorescent beads, as fluorophores can alter the light scatter properties of beads. Light scatter calibration should preferably be performed on samples acquired by SSC-trigger threshold. This allowed for confident determination of the limit of light scatter sensitivity. If a fluorescent trigger is used and the SSC intensities are below those that could be acquired with a SSC-trigger threshold it is critical to demonstrate that the models retain accuracy in the region they are being extrapolated to. For fluorescence calibration, beads with matching fluorophore to the biological particle were used. This was essential for the fluorescence intensities to be comparable between instruments with differing optical configurations, such as filter sets and laser wavelengths for excitation^24^. In summary, the methods employed here compellingly show that standardized data reporting of fluorescence and light scatter is achievable for small particle FCM data when it is converted to standard units of MESF and diameter.

It is important to stress that the primary purpose of light scatter calibration is not to determine the absolute size of biological particles, but to demonstrate a means of comparing data in standardized units with a consistent population. MV-M-sfGFP was chosen as a proof-of-principle sample due to it being a well-defined, homogeneous population. The assumed effective RI of the virus, 1.45^18^, significantly impacts the particle diameter predicted by FCM_PASS_, which in this case is congruent with published EM measurements of the virus. Validation of these methodologies for the extracellular vesicle (EV) field are required, and will be more challenging due to the heterogeneity of EV populations, with the majority of particles likely below the triggering threshold with a log-normal distribution^21^. This will mean that normalization of the light scatter and fluorescence data between instruments will heavily rely upon accurate quantitation of the concentration. This is challenging as variations can occur in concentration with identical samples on the same platform, as we have shown in this pilot study. Further testing of fluidic stability and acquisition time for their effect on recorded concentration variation with an easily detectable, homogeneous population should therefore be a priority for laboratories attempting to use concentration measurements and would be the next step in this line of work.

Ultimately, calibration is critical to allow for meaningful biological conclusions and allow reproduction of work. The use of calibration can also aid researchers in identifying optimal assays and equipment for their work. Literature is beginning to emerge where fluorescence calibration is implemented, however, small particle studies and standardization efforts have predominantly used light scatter calibration as a normalization method^11,25–30^. While light scatter calibration has great utility, fluorescence calibration should be seen as a priority in any study utilizing fluorescence staining, whether it is immunophenotyping or fluorescent dyes. This is due to the general convention of reporting only the counts of fluorescent events instead of total detected counts of all events. To this end, a follow up standardization study is required that utilizes both fluorescence and light scatter calibration to compare instrument sensitivities and further validate the findings of this pilot study.

While the need for calibration is essential for the field to progress, its use alone does not ensure quality of research or that instrument settings are optimal. The use of optimal experimental design with controls such as serial dilution to ensure single particle detection, buffer with reagent controls to demonstrate lack of artefacts, and isotype or negative staining controls to show specificity are all required^15^. A consensus for reporting of the methods and experimental design needs to be utilized for researchers to make easy comparisons between studies and allow for reproduction of data. This effort is actively being undertaken by the International Society of Extracellular Vesicles, International Society for Advancement of Cytometry, and International Society for Thrombosis and Haemostasis flow cytometry working group flow cytometry working group who recently laid out a standard reporting framework (MIFlowCyt-EV) and minimal experiment requirement^15^.

---

**Table 1 – Summary of scatter and fluorescence gain/voltage settings used for detection of anti-GFP PE labeled MV-M-sfGFP.**

**Table 2 – Summary of statistics obtained for detection of anti-GFP PE labeled MV-M-sfGFP**. Light scatter and fluorescence statistics were calculated using a gate from 50-400 PE MESF units, with the 50^th^ percentile (5^th^, 95^th^ percentile) shown. Count and concentration statistics were calculated using a 50-400 PE MESF gate with a comparison gate at 50-400 PE MESF and 118-180 nm to account for the trigger threshold of the LSR Fortessa which appears to cut of a small portion of the MV-M-sfGFP population.

**Supplementary Table 1 – Completed MIFlowCyt framework**

**Supplementary Table 2 – Completed MIFlowCyt-EV framework**

**Supplementary Table 3 – Light scatter modelling outputs of CytoFLEX S.** The modelling input settings for the FCM_PASS_ software for flow cytometer calibration are listed. Bead diameter, diameter CV, and refractive index (RI) inputs are based on manufacturer provided information.

**Supplementary Table 4 – Light scatter modelling outputs of LSR Fortessa.** The modelling input settings for the FCM_PASS_ software for flow cytometer calibration are listed. Bead diameter, diameter CV, and refractive index (RI) inputs are based on manufacturer provided information.

**Supplementary Table 5 – Summary of cross-calibrated PE MESF values from the CytoFLEX S and LSR Fortessa.**

**Supplementary Figure 1 – Full dataset demonstrating the comparability of uncalibrated and calibrated data across flow cytometers. A-F)** The side scatter versus anti-GFP-PE fluorescence of MV-M-sfGFP are shown on the LSR Fortessa and CytoFLEX S, respectively. A comparison gate is drawn from 4 x 10^3^ to 6 x 10^4^ SSC-H and 1 x 10^3^ to 3 x 10^4^ PE-A. **G-L)** The scatter cross section versus Anti-GFP-PE (MESF) of MV-M-sfGFP are shown on the LSR Fortessa and CytoFLEX S, respectively. A comparison gate is drawn from 1 x 10^3^ to 1 x 10^4^ nm^2^ and 20 to 400 PE MESF. **M-R)** The diameter (nm) versus Anti-GFP-PE (MESF) of MV-M-sfGFP are shown on the LSR Fortessa and CytoFLEX S, respectively. A comparison gate is drawn from 100 to 180 nm to and 20 to 400 PE MESF (black). A second comparison gate is drawn from 118 to 180 nm to and 50 to 400 PE MESF (red dotted). **S-X)** The diameter (nm) versus Anti-GFP-PE (MESF nm^−2^) of MV-M-sfGFP are shown on the LSR Fortessa and CytoFLEX S, respectively. A comparison gate is drawn from 100 to 180 nm to and 5 x 10^−4^ to 8 x 10^−3^ PE MESF nm^−2^ (black). Flow cytometer platform and acquisition setting number are outlined at the top. Information on acquisition settings can be found in **Table 1**.

**Supplementary Figure 2 - Light scatter modelling outputs of CytoFLEX S.** The model fit (A) depicts the predicted versus acquire scatter for each bead. Refractive indices (RIs) are differentiated by color. The predicted collection angle (B) shows the collection angle used to create the scatter-diameter model (C). The median scatter intensity of anti-GFP-PE labelled MV-M-sfGFP assuming its diameter is 124 nm is shown in blue. The cytometer configuration and data acquired from bead populations used to model the CytoFLEX S and produce these figures are summarized in **Supplementary Information 1**.

**Supplementary Figure 3 - Light scatter modelling outputs of LSR Fortessa.** The model fit (A) depicts the predicted versus acquire scatter for each bead. Refractive indices (RIs) are differentiated by color. The predicted collection angle (B) shows the collection angle used to create the scatter-diameter model (C). The median scatter intensity of anti-GFP-PE labelled MV-M-sfGFP assuming its diameter is 124 nm is shown in blue. The cytometer configuration and data acquired from bead populations used to model the LSR Fortessa and produce these figures are summarized in **Supplementary Information 2**.

**Supplementary Figure 4 – Control dataset across flow cytometers and detector settings.** Control data for the CytoFLES S (**A-K**) and LSR Fortessa (**L-V**) are represented at detector settings outlined in **Table 1**. All plots are shown as scattering cross section vs. PE MESF. A gate is drawn at 20 MESF units (red) and 50 MESF units (black) for all plots.

