## Supplementary material for "Fluorescence and light scatter calibration allow comparisons of small particle data in standard units across different flow cytometry platforms and detector settings": Table 1

|  | **CytoFLEX S (Gain)** | | | **LSR Fortessa (Voltage)** | | |
| --- | --- | --- | --- | --- | --- | --- |
| **Setting** | 1 | 2 | 3 | 1 | 2 | 3 |
| **SSC Channel** | 125 | 180 | 250 | 410 | 410 | 410 |
| **GFP Channel** | 3000 | 3000 | 3000 | 540 | 540 | 540 |
| **PE Channel** | 300 | 1000 | 2000 | 463 | 605 | 650 |
| **Trigger** | VSSC | VSSC | VSSC | SSC | SSC | SSC |
| **Threshold** | 1400 | 1500 | 2000 | 200 | 200 | 200 |
