## Supplementary material for "Fluorescence and light scatter calibration allow comparisons of small particle data in standard units across different flow cytometry platforms and detector settings": Table 2

|  | **CytoFLEX S** | | | **LSR Fortessa** | | |
| --- | --- | --- | --- | --- | --- | --- |
| **Setting** | **1** | **2** | **3** | **1** | **2** | **3** |
| **Light Scatter**  (a.u.) | **6542** (4790, 10264) | **9349** (6722, 13430) | **13121** (9425, 19751) | **348**  (240, 620) | **348** (240, 620) | **347** (240, 632) |
| **Scatter Cross Section**  (nm^2^) | **1894** (1387, 2981) | **1889** (1358, 2723) | **1903** (1367, 2889) | **1326** (915, 2358) | **1326** (915, 2349 | **1323** (915, 2409 |
| **Diameter**  (nm) | **120.7** (113, 133) | **120.6**  (113, 130) | **120.8** (113, 132) | **130.3** (121, 146) | **130.3** (121, 146) | **130.2** (121, 146) |
| **PE**  (a.u.) | **2117** (1232, 3368) | **8925** (4824, 12432) | **17604** (9630, 25227) | **111** (64, 175) | **847** (508, 1283) | **1454** (894, 2204) |
| **PE**  (MESF) | **99** (58, 156) | **128** (70, 178) | **128** (70, 182) | **110** (63, 173) | **108** (64, 164) | **103** (62, 157) |
| **PE**  (x 10^-3^ MESF nm^-2^) | **2.16** (0.133, 0.310) | **2.80** (0.160, 0.362) | **2.78** (0.162, 0.366) | **2.06** (0.120, 0.300) | **2.02** (0.122, 0.285) | **1.93** (0.119, 0.273) |
| **Count x10^4^** (50-400 PE MESF) | **1.38** | **1.85** | **1.68** | **2.55** | **2.05** | **2.38** |
| **Count x10^4^** (50-400 PE MESF, 118-180 nm) | **1.12** | **1.49** | **1.37** | **2.55** | **2.05** | **2.38** |
| **Concentration x10^6^ mL^-1^** (50**-**400 PE MESF) | **1.46** | **1.75** | **1.69** | **1.25** | **1.19** | **1.24** |
| **Concentration x10^6^ mL^-1^**  (50-400 PE MESF, 118-180 nm) | **1.18** | **1.41** | **1.38** | **1.25** | **1.19** | **1.24** |
