## Supplementary Table 1 for "Fluorescence and light scatter calibration allow comparisons of small particle data in standard units across different flow cytometry platforms and detector settings"

**Cytometry Part A**

**Author Checklist: MIFlowCyt-Compliant Items**

| **Requirement** | **Please Include Requested Information** |
| --- | --- |
| 1.1. Purpose | The purpose of this technical note was to demonstrate that scatter and fluorescence calibration for standardized data reporting. |
| 1.2. Keywords | Scatter Calibration, Fluorescence Calibration, Validation materials, Calibration materials, Reference Particles, MESF |
| 1.3. Experiment variables | Parameters detected on both cytometers: side-scatter (SSC), sfGFP, phycoerythrin (PE) |
| 1.4. Organization name and address | University of Ottawa, 451 Smyth Rd. Roger Guindon Hall, K1H8M5, Ottawa, ON. Canada |
| 1.5. Primary contact name and email address | Joshua Welsh,  Vera Tang, |
| 1.6. Date or time period of experiment | May - August 2019 |
| 1.7. Conclusions | Scatter and fluorescence calibration allowed for direct comparison of data collected using the same biological sample from two different flow cytometry platforms |
| 1.8. Quality control measures | CS&T was performed through BD FACSDiva for the LSRFortessa. QC was performed using CytExperts on the CytoFLEX S. Flow rate was calibrated on both cytometers as described in Methods section. Scatter and fluorescence calibration was performed as described in Methods section. Reagent only controls (anti-GFP PE only) and buffer only controls were included. |
| 2.1.1.1. (2.1.2.1., 2.1.3.1.) Sample description | NIST traceable polystyrene and silica beads, Quantibrite PE beads – see Materials and Methods section for description. MV-M-sfGFP – see below for description. |
| 2.1.1.2. Biological sample source description | MV-M-sfGFP is Moloney strain murine leukemia virus (MLV) with superfolder GFP inserted in fusion with the envelope glycoprotein. |
| 2.1.1.3. Biological sample source organism description | MV-M-sfGFP is produced by chronic infection of NIH3T3 mouse fibroblast cells |
| 2.1.2.2. Environmental sample location | NA |
| 2.3. Sample treatment description | MV-M-sfGFP was labeled with anti-GFP-PE. All samples (beads and virus) were diluted and analyzed in phosphate buffered saline. |
| 2.4. Fluorescence reagent(s) description | MV-M-sfGFP expresses sfGFP. Anti-GFP conjugated with phycoerythrin (PE) was used to label MV-M-sfGFP. Other fluorescent reagents include Quantibrite PE MESF beads, which contain PE. |
| 3.1. Instrument manufacturer | Beckman Coulter, BD Biosciences |
| 3.2. Instrument model | Beckman Coulter CytoFLEX S, BD LSR Fortessa |
| 3.3. Instrument configuration and settings | CytoFLEX S (100mW 405nm, 50mW 488nm, 50mW 561nm, 50mW 640nm)  405SSC – 405/10, 488-530/30, 561-586/15  LSRFortessa (50mW 405nm, 50mW 488nm, 50mW 561nm, 50mW 640nm)  488SSC – 488/10, 488-525/40, 561-585/15 |
| 4.1. List-mode data files  *We recommend all authors to submit their data files to [http://flowrepository.org](http://flowrepository.org/) and to make them available for the peer-review process. If you have done so, please let us know by inserting the following codes (replace the red text):  1) The link for peer-review process:  http://flowrepository.org/id/RvFrxxxxxx (copy and paste the code). This link will only be shared with reviewers of your manuscript.  2) The repository identifier:  http://flowrepository.org/id/FR-FCM-xxxx (copy and paste the code). This link will be made publicly accessible after the paper is published. | https://jones-lab-nanopass.s3.amazonaws.com/Paper+Data/2020/Cyto+Part+A%2C+Technical+Note/Archive.zip |
| 4.2. Compensation description | No compensation was performed |
| 4.3. Data transformation details | See MIFlowCytEV Framework for detailed description of data transformation of SSC to diameter and fluorescence in arbitrary units to MESF (<http://www.evflowcytometry.org/links/>) |
| 4.4.1. Gate description | See Methods section and Figure 1 of manuscript |
| 4.4.2. Gate statistics | See Methods section and Figure 2 of manuscript |
| 4.4.3. Gate boundaries | See Methods section and Figures 1 & 2 of manuscript |

**Notes**

Feel free to use more space than allocated.

You can embed graphics/figures in this document, if needed.

Please make sure to save the document in Microsoft Word version 2003 or older, before uploading to ScholarOne Manuscripts. When uploading this checklist to ScholarOne Manuscripts, please choose the “Supplementary Material for Review” category.

Please note that if your paper is accepted, the checklist will be published as an Online Supporting Information.

For any questions, please contact the Cytometry Part A editorial office at.
