## Supplementary Table 2 for "Fluorescence and light scatter calibration allow comparisons of small particle data in standard units across different flow cytometry platforms and detector settings"

| **Framework Criteria** | **What to report** | **Please complete each criterion** |
| --- | --- | --- |
| 1.1 Preanalytical variables conforming to MISEV guidelines. | Preanalytical variables relating to EV sample including source, collection, isolation, storage, and any others relevant and available in the performed study. | MLVsfGFP and MLV no GFP was acquired from Viroflow Technologies. Storage: 4C. MLVsfGFP and MLV no GFP are produced from chronically infected NIH3T3 cells, isolated from infected cell supernatants. Virus is deactivated with formalin and lyophilized. Product information sheet can be found online: https://static1.squarespace.com/static/5bc8a4d2c46f6d604a42e518/t/5d8384d084be497be56a7def/1568900304843/ProductInformationSheet+v3.3.pdf |
| 1.2 Experimental design according to MIFlowCyt guidelines. | EV-FC manuscripts should provide a brief description of the experimental aim, keywords, and variables for the performed FC experiment(s) using MIFlowCyt checklist criteria: 1.1, 1.2, and 1.3, respectively | See MIFlowCyt Checklist sections 1.1, 1.2, 1.3 |
| 2.1 Sample staining details | State any steps relating to the staining of samples. Along with the method used for staining, provide relevant reagent descriptions as listed in MIFlowCyt guidelines (Section 2.4 Fluorescence Reagent(s) Descriptions). | See MIFlowCyt Checklist sections 2.4. MV-M-sfGFP (ViroFlow Technologies, Canada) was re-constituted as per manufacturer's instructions. The concentration of virus particles was adjusted to 1x109 particles mL-1 according to the concentration provided by the manufacturer and labeled with anti-GFP PE antibody (Clone FM264G, BioLegend) at 0.4 µg mL-1 for a minimum of 30 minutes at room temperature, protected from light. |
| 2.2 Sample washing details | State any steps relating to the washing of samples. | No washing steps were performed. |
| 2.3 Sample dilution details | All methods and steps relating to sample dilution. | MV-M-sfGFP (ViroFlow Technologies, Canada) was re-constituted as per manufacturer's instructions. The concentration of virus particles was adjusted to 1x109 particles mL-1 according to the concentration provided by the manufacturer for antibody labeling. Antibody labeled virus was diluted to ~1x106 particles mL-1 immediately prior to analysis by flow cytometry. Antibody alone and unstained virus samples were similarly prepared. All dilutions were made using 0.1 µm filtered PBS (PBS 1x, no Ca2+, no Mg2+, Wisent). |
| 3.1 Buffer alone controls. | State whether a buffer-only control was analyzed at the same settings and during the same experiment as the samples of interest. If utilized it is recommended that all samples be recorded for a consistent set period of time e.g. 5 minutes, rather than stopping analysis at a set recorded event count e.g. 100,000 events. This allows comparisons of total particle counts between controls and samples. | A buffer only control (PBS) was analyzed. |
| 3.2 Buffer with reagent controls. | State whether a buffer with reagent control was analyzed at the same settings, same concentrations, and during the same experiment as the samples of interest. If used state what the results were. | A buffer with antibody only control (PBS-aGFP PE) was analyzed at the same settings during the same experiment as the samples of interest, at the same concentration, acquired for the same duration. |
| 3.3 Unstained controls. | State whether unstained control samples were analyzed at the same settings and during the same experiment as stained samples. If used, state what the results were, preferably in standard units. | An unstained MLVsfGFP (virus) sample was analyzed. 1 PE MESF unit for unstained virus on both instruments |
| 3.4 Isotype controls. | The use of isotype controls is applicable to immunofluorescence labelling only. State whether isotype controls were analyzed at the same settings and during the same experiment as stained samples. If utilized, state which antibody they are matched to, the concentration used, and what the results were (Section 4.2, 4.3, 4.4). Due to conjugation differences between manufacturers if should be stated if the isotype controls are from the same manufacturer as the matched antibodies. | Isotype control was not used. MLV with noGFP was instead used as a negative staining control |
| 3.5 Single-stained controls. | State whether single-stained controls were included. If used state whether the single-stained controls were recorded using the same settings, dilutions, and during the same experiment as stained samples and state what the results were, preferably in standard units (Section 4.2, 4.3, 4.4). | MLVsfGFP was used as a single stain control |
| 3.6 Procedural controls. | State whether procedural controls were included. If used, state the procedure and if the procedural controls were acquired at the same settings and during the same experiment as stained samples. | Not applicable |
| 3.7 Serial dilutions. | State whether serial dilutions were performed on samples and note the dilution range and manner of testing. The fluorescence and/or scatter signal intensity would ideally be reported in standard units (see Section 4.3, 4.4) but arbitrary units can also be used. This data is best reported by plotting the recorded number events/concentration over a set period of time at different sample dilution. The median fluorescence intensity at each of the dilutions should also ideally be plotted on the same or a separate plot. | The detected virus is a monomodal population above the detection sensitivity of the instrument. The sample was analysed at 1x10^6 particle per mL based on manufacturer's measurements using flow cytometry and nanoparticle tracking analysis. |
| 3.8. Detergent treated EV-samples | State whether samples were detergent treated to assess lability. If utilized, state what detergent was used, the end concentration of the detergent, and what the results were of the lysis. | Samples were not treated with detergent |
| 4.1 Trigger Channel(s) and Threshold(s). | The trigger channel(s) and threshold(s) used for event detection. Preferably, the fluorescence calibration (Section 4.3) and/or scatter calibration (Section 4.4) should be used in order to report the trigger channel(s) and threshold(s) in standardized units. | Set 1, 2, 4 = 200 au (~800nm^2) Fortessa SSC, Set 1, 2, 3 = 1500, 2000, 1400 (~300 nm^2) Cytoflex VSSC. |
| 4.2 Flow Rate / Volumetric quantification. | State if the flow rate was quantified/validated and if so, report the result and how they were obtained. | The flow rate on the CytoFlex was calibrated with samples measured at 15 µl/min. The flow rate on the LSR Fortessa was calculated by cross-calibrating the concentration of 200 nm fluorescent polystyrene spike-in beads (Green FluoSpheres, Cat# F8848, ThermoFisher Scientific, USA) using the mean concentration of beads acquired using the CytoFLEX S after calibration of the fluidic system at a low flow rate. The LSR Fortessa flow rate was calculated to be ~18 µL/min. |
| 4.3 Fluorescence Calibration. | State whether fluorescence calibration was implemented, and if so, report the materials and methods used, catalogue numbers, lot numbers, and supplied reference units for the standards. Fluorescence parameters may be reported in standardized units of MESF, ERF, or ABC beads. The type of regression used, and the resulting scatter plot of arbitrary data vs standard data for the reference particles should be supplied. | Fluorescence calibration was performed using PE MESF beads (QuantiBrite PE, Cat# 340495, Lot 73318, BD Biosciences, USA) at a voltage/gain that allowed for the brightest bead to be within the range of detection of the instrument. Median PE was gated using FlowJo. Fluorescence calibration was performed using FCMPASS software. These PE MESF values were used to cross-calibrated 8-peak rainbow bead (Cat# RCP-30-5A, Lot AF01, Spherotech, USA) data acquired at the same settings as the PE MESF beads. The 8-peak rainbow beads were then used to calibrate the PE intensity scales at different acquisition settings. This is a cost effective method demonstrating the cross calibration of different types of fluorescence calibration beads, where the 8-peak rainbow beads will also have more populations on-scale at the high voltage/gain settings. |
| 4.4 Light Scatter Calibration. | State whether and how light scatter calibration was implemented. Light scatter parameters may be reported in standardized units of nm2, along with information required to reproduce the model. | Light scatter calibration was performed using 81, 100, 152, 203, 269, 303, 345, 401, 453, 568, 600 nm polystyrene NIST-traceable beads (ThermoFisher Scientific, USA) and 480 and 730 nm silica NIST-traceable beads (ThermoFisher Scientific, USA). Median SSC intensity (488 nm SSC on LSR Fortessa, 405 nm SSC on CytoFLEX S) were gated using FlowJo (v10.5.3, USA). Mie modeling and subsequent conversion of light scatter intensity to diameter was performed using FCMPASS software (http://nanopass.ccr.cancer.gov)[welsh et al, 2019, cyto part a]. Model input settings including refractive indices, bead information, and statistics can be found in Supplementary Information 1 and 2, with model outputs shown in Supplementary Figure 1 and 2. The collection half-angle refers to the angle of light being collected around the particle within the sheath of the instrument. |
| 5.1 EV diameter/surface area/volume approximation. | State whether and how EV diameter, surface area, and/or volume has been calculated using FC measurements. | Calibration of light scatter to a diameter in nanometers resulted in median values that ranged from just 121 to 130 nm for MLVsfGFP. |
| 5.2 EV refractive index approximation. | State whether the EV refractive index has been approximated and how this was done. | Virus refractive index was approximated to be 1.45 at 488 nm assume the size of the virus was ~124 nm based on the published literature. |
| 5.3 EV epitope number approximation. | State whether EV epitope number has been approximated, and if so, how it was approximated. | Median epitope number ~99-128, assuming no steric hindrance, one PE molecule per antibody, one antibody per GFP molecule, and accurate regression |
| 6.1 Completion of MIFlowCyt checklist. | Complete MIFlowCyt checklist criteria 1 to 4 using the MIFlowCyt guidelines. | See attached MIFlowCyt Checklist |
| 6.2 Calibrated channel detection range | If fluorescence or scatter calibration has been carried out, authors should state whether the upper and lower limits of a calibrated detection channel were calculated in standardized units. This can be done by converting the arbitrary unit scale to a calibrated scaled, as discussed in Section 4.3 and 4.4, and providing the highest unit on this scale and the lowest detectable unit above the unstained population. The lowest unit at which a population is deemed ‘positive’ can be determined a variety of ways, including reporting the 99th percentile measurement unit of the unstained population for fluorescence. The chosen method for determining at what unit an event was deemed positive should be clearly outlined. | The PE intensity of anti-GFP PE labeled MV-M-sfGFP from the LSR Fortessa and CytoFLEX S ranged in arbitrary units of intensity from 110 to 17322 using three different gain/voltage settings as defined in the methods section, Figure 2A and 2B. Upon calibration of fluorescence intensity to PE MESF units, values ranged from just 89 to 127 PE MESF. Data irrespective of instrument or instrument settings were comparable. hree scatter settings were tested on the CytoFLEX S and LSR Fortessa (defined in methods section). Light scatter intensity for SSC across the LSR Fortessa and CytoFLEX S ranged from 346 to 13014 arbitrary units, Figure 2C and 2D. Calibration of light scatter to diameter in nanometers resulted in values that ranged from just 123 to 126 nm. |
| 6.3 EV number/concentration. | State whether EV number/concentration has been reported. If calculated, it is preferable to report EV number/concentration in a standardized manner, stating the number/concentration between a set detection range. | Virus concentration on the CytoFLEX S and LSR Fortessa was calculated using 200 nm fluorescent polystyrene spike-in beads (Green FluoSpheres, Cat# F8848, ThermoFisher Scientific, USA) using the mean concentration of beads acquired using the CytoFLEX S after calibration of the fluidic system at a low flow rate. |
| 6.4 EV brightness. | When applicable, state the method by which the brightness of EVs is reported in standardized units of scatter and/or fluorescence. | EV brightness was shown on dot plots with standard units of fluorescence (PE MESF) and light scatter (nm). See sections 4.3 and 4.4 |
| 7.1. Sharing of data to a public repository. | Provide a link to the experimental data in a public data repository. | <https://jones-lab-nanopass.s3.amazonaws.com/Paper+Data/2020/Cyto+Part+A%2C+Technical+Note/Archive.zip> |
