## Supplementary Table 3 for "Fluorescence and light scatter calibration allow comparisons of small particle data in standard units across different flow cytometry platforms and detector settings"

| Flow Cytometer Modelling Settings |  |  |  |  |  |  |  |
| --- | --- | --- | --- | --- | --- | --- | --- |
| FCMPASS software version | 2.16 |  |  |  |  |  |  |
| Flow cytometer model | CytoFLEX |  |  |  |  |  |  |
| Acquisition Date | 2019-07-06 |  |  |  |  |  |  |
| Flow cytometer illumination wavelength | 405 |  |  |  |  |  |  |
| Suspension Medium Refractive Index | 1.3427 |  |  |  |  |  |  |
| Flow cytometer collection angle approximation | On |  |  |  |  |  |  |
| Flow cytometer closest fitting collection geometry | Circle |  |  |  |  |  |  |
| Flow cytometer Collection Angles | T1=90, P1=90, E1=53.2 |  |  |  |  |  |  |
| Flow cytometer Calibration Factor | 0.209211542 |  |  |  |  |  |  |
| Vesicle Modelling Settings |  |  |  |  |  |  |  |
| Vesicle Average Cytosol Refractive Index | 1.3859 |  |  |  |  |  |  |
| Vesicle Upper Cytosol Refractive Index | 1.406 |  |  |  |  |  |  |
| Vesicle Lower Cytosol Refractive Index | 1.3658 |  |  |  |  |  |  |
| Vesicle Membrane Refractive Index | 1.4863 |  |  |  |  |  |  |
| Vesicle Membrane Thickness (nm) | 10 |  |  |  |  |  |  |
| Bead Modelling Settings |  |  |  | Set 1 | Set 2 | Set 3 |  |
| Diameter (nm) | Diameter (%CV) | Bead RI | SSC (%CV) | SSC Value (a.u.) | | |  |
| 81 | 11.7 | 1.6253 | 34.8 | 5119 | 7347 | 10241 | Reference |
| 100 | 6.8 | 1.6253 | 19.6 | 15312 | 21975 | 30632 | Reference |
| 152 | 3.3 | 1.6253 | 7.77 | 100915 | 144828 | 201883 | Reference |
| 203 | 2.6 | 1.6253 | 7.19 | 270736 | 388547 | 541615 | Reference |
| 269 | 1.6 | 1.6253 | 5.06 | 608980 | 873977 | 1218280 | Reference |
| 303 | 1.6 | 1.6253 | 2.21 | 772546 | 1108718 | 1545498 | Reference |
| 345 | 1.9 | 1.6253 | 2.99 | 902670 | 1295466 | 1805815 | Reference |
| 401 | 1.3 | 1.6253 | 2.62 | 1003344 | 1439947 | 2007214 | Reference |
| 453 | 1.7 | 1.6253 | 2.5 | 1207286 | 1732635 | 2415207 | Reference |
| 508 | 1.7 | 1.6253 | 4.11 | 1672624 | 2400463 | 3346126 | Reference |
| 600 | 1.7 | 1.6253 | 5.7 | 3135952 | 4500556 | 6273550 | Reference |
| 480 | 4.2 | 1.4611 | 5.38 | 174778 | 250832 | 349647 | Reference |
| 730 | 4.1 | 1.4611 | 13.6 | 699125 | 1003348 | 1398617 | Reference |
| 0 | 0 | 1.45 | 0 |  | 0 |  | Curve |
| 124 | 5.5 | 0 | 36 |  | 8236 |  | Data Point |
