## Supplementary Table 4 for "Fluorescence and light scatter calibration allow comparisons of small particle data in standard units across different flow cytometry platforms and detector settings"

| Flow Cytometer Modelling Settings |  |  |  |  |  |
| --- | --- | --- | --- | --- | --- |
| FCMPASS software version | 2.17 |  |  |  |  |
| Flow cytometer model | Fortessa |  |  |  |  |
| Acquisition Date | 2019-07-06 |  |  |  |  |
| Flow cytometer illumination wavelength | 488 |  |  |  |  |
| Suspension Medium Refractive Index | 1.337 |  |  |  |  |
| Flow cytometer collection angle approximation | On |  |  |  |  |
| Flow cytometer closest fitting collection geometry | Circle |  |  |  |  |
| Flow cytometer Collection Angles | T1=90, P1=90, E1=46.6 |  |  |  |  |
| Flow cytometer Calibration Factor | 3.169105636 |  |  |  |  |
| Vesicle Modelling Settings |  |  |  |  |  |
| Vesicle Average Cytosol Refractive Index | 1.38 |  |  |  |  |
| Vesicle Upper Cytosol Refractive Index | 1.4 |  |  |  |  |
| Vesicle Lower Cytosol Refractive Index | 1.36 |  |  |  |  |
| Vesicle Membrane Refractive Index | 1.48 |  |  |  |  |
| Vesicle Membrane Thickness (nm) | 10 |  |  |  |  |
| Bead Modelling Settings |  |  |  |  |  |
| Diameter (nm) | Diameter (%CV) | Bead RI | SSC (%CV) | SSC Value (a.u.) | |
| 152 | 3.3 | 1.6039 | 11.99 | 5203 | Reference |
| 203 | 2.6 | 1.6039 | 7.19 | 14066 | Reference |
| 269 | 1.6 | 1.6039 | 5.06 | 30280 | Reference |
| 303 | 1.6 | 1.6039 | 2.21 | 39315 | Reference |
| 345 | 1.9 | 1.6039 | 2.99 | 46575 | Reference |
| 401 | 1.3 | 1.6039 | 2.62 | 55947 | Reference |
| 453 | 1.7 | 1.6039 | 2.5 | 60137 | Reference |
| 600 | 1.7 | 1.6039 | 5.7 | 136090 | Reference |
| 480 | 4.2 | 1.4546 | 5.38 | 10479 | Reference |
| 730 | 4.1 | 1.4546 | 13.6 | 38238 | Reference |
| 0 | 0 | 1.4453 | 0 | 0 | Curve |
| 124 | 5.5 | 0 | 0 | 345.8 | Data Point |
