## Supplementary Table 5 for "Fluorescence and light scatter calibration allow comparisons of small particle data in standard units across different flow cytometry platforms and detector settings"

| **Population** | **CytoFLEX S** | **LSR Fortessa** | **Difference (%)** |
| --- | --- | --- | --- |
| **1** | 8 | 8 | 0.00 |
| **2** | 200 | 195 | 2.56 |
| **3** | 555 | 552 | 0.54 |
| **4** | 1678 | 1701 | 1.35 |
| **5** | 4289 | 4395 | 2.41 |
| **6** | 12181 | 12574 | 3.13 |
| **7** | 37095 | 38272 | 3.08 |
| **8** | 92487 | 92666 | 0.19 |
