## Supplementary figures and images for "Fluorescence and light scatter calibration allow comparisons of small particle data in standard units across different flow cytometry platforms and detector settings"

### Supplementary Figure 1

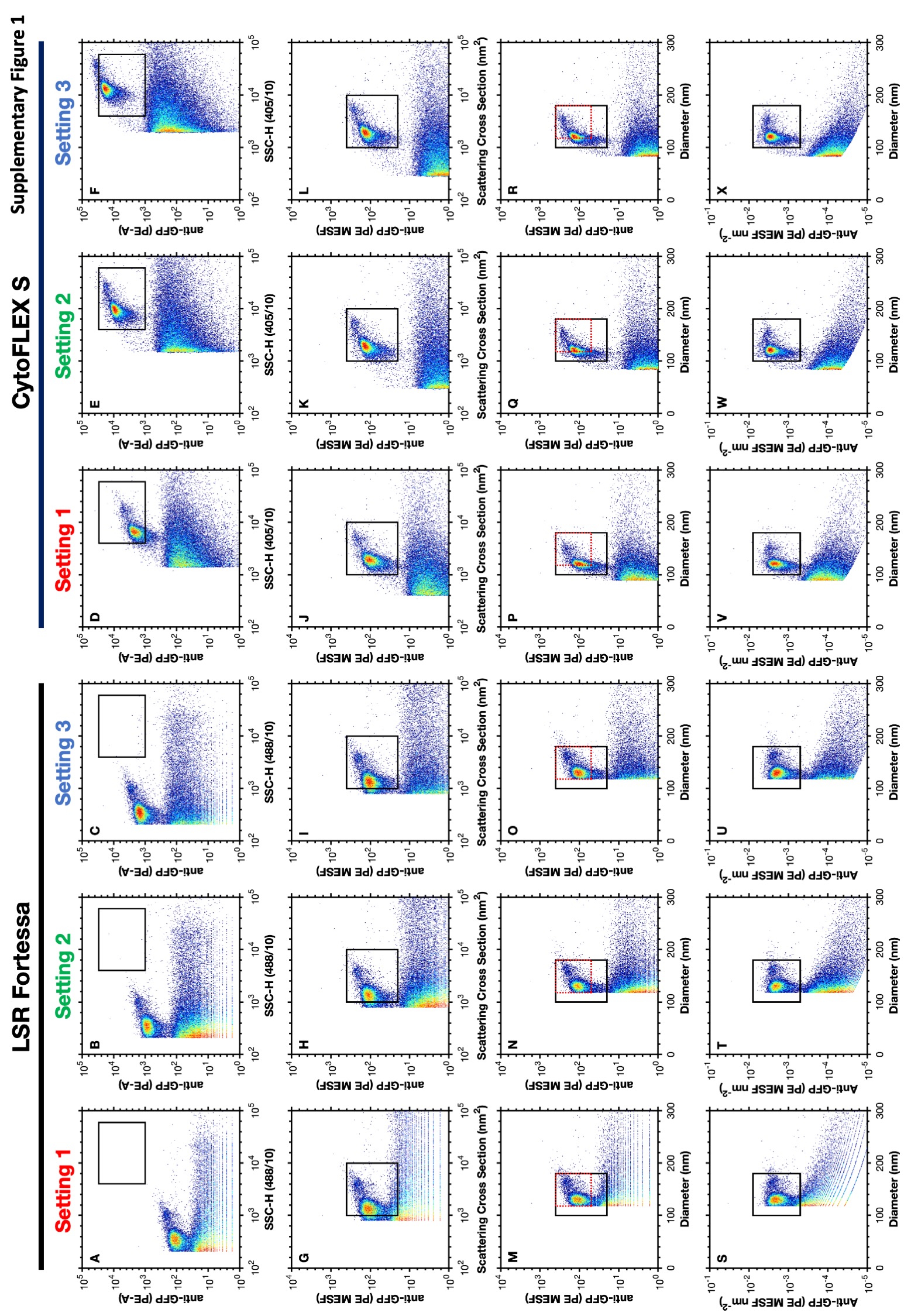

### Supplementary Figure 2

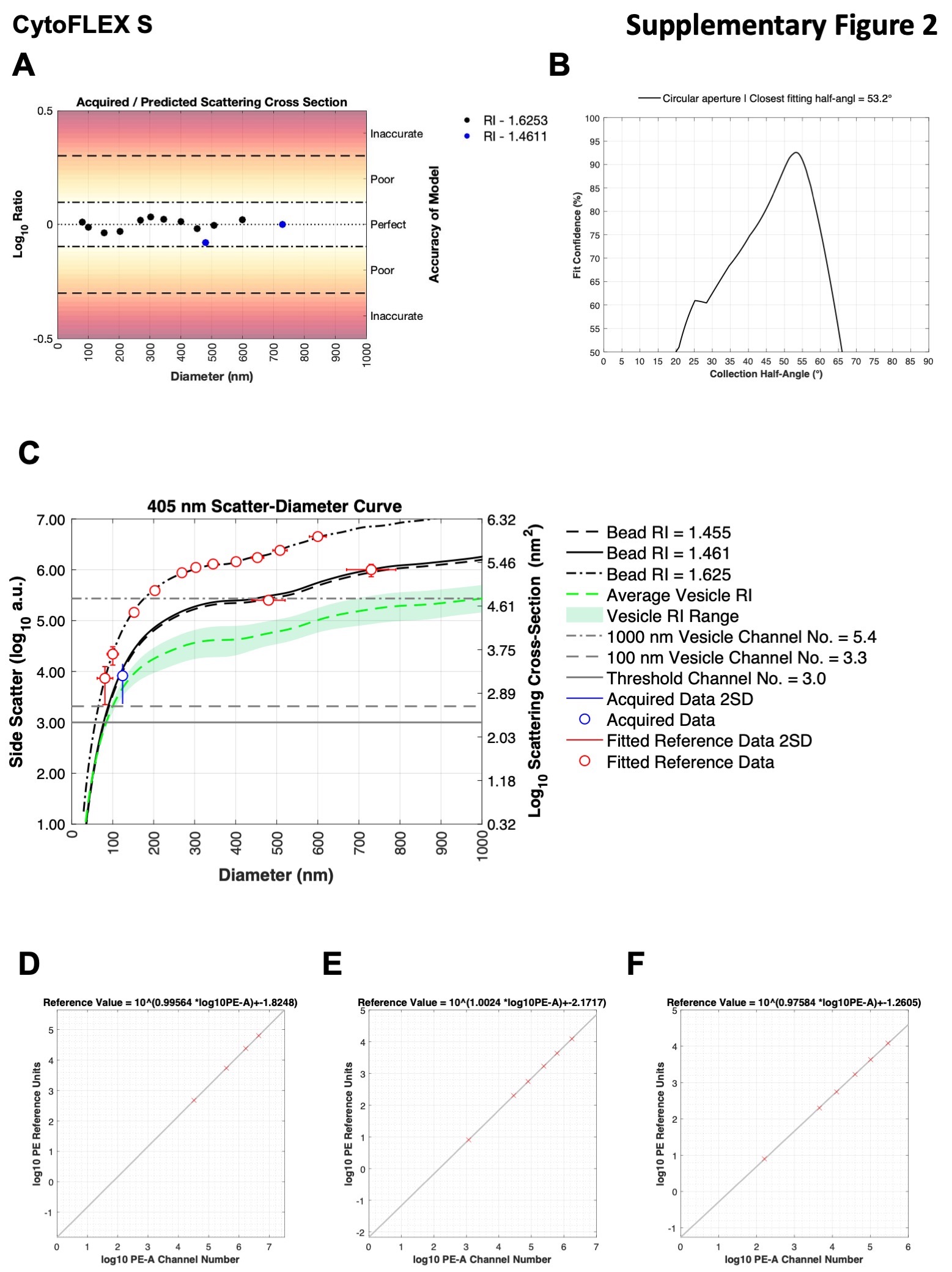

### Supplementary Figure 3

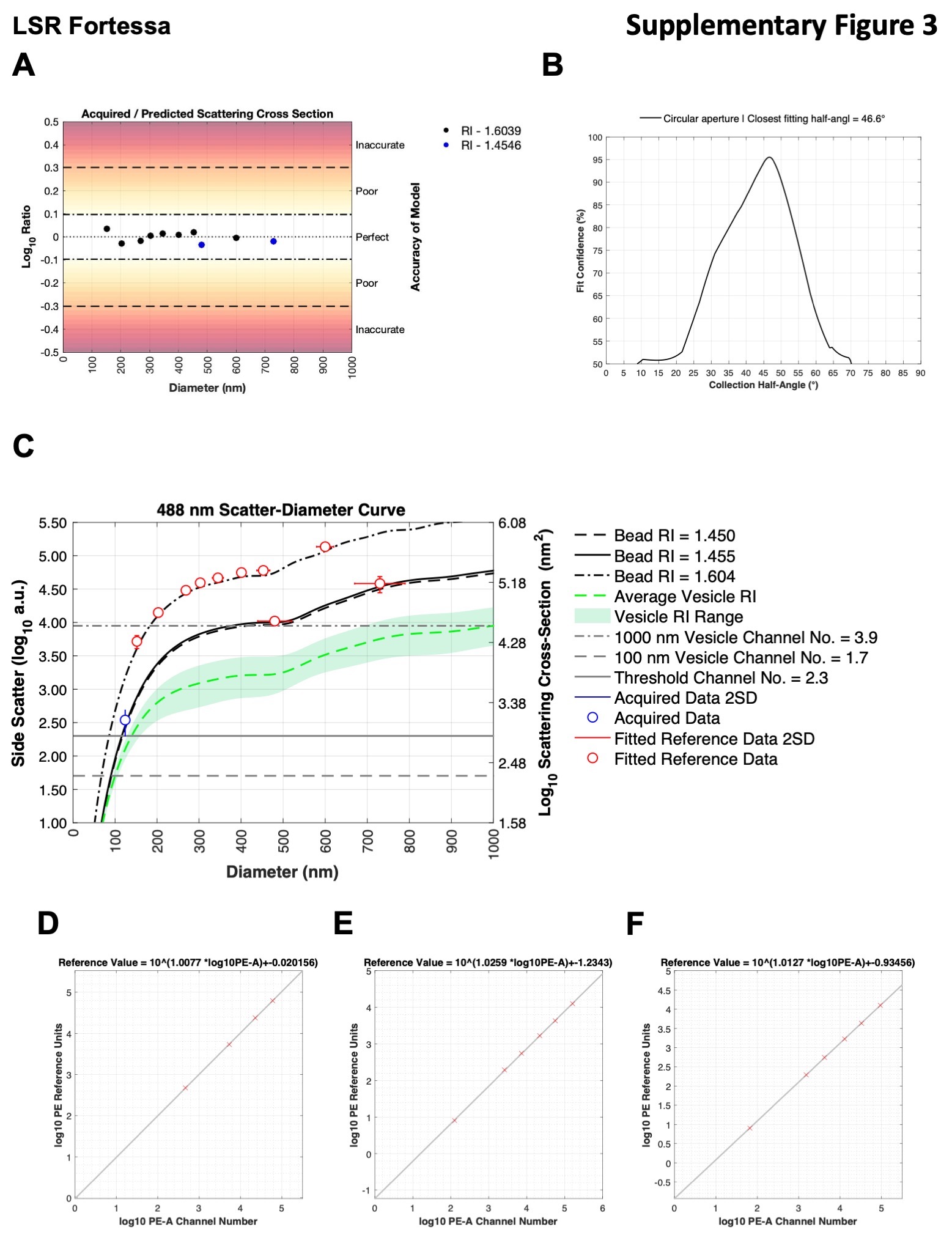

### Supplementary Figure 4

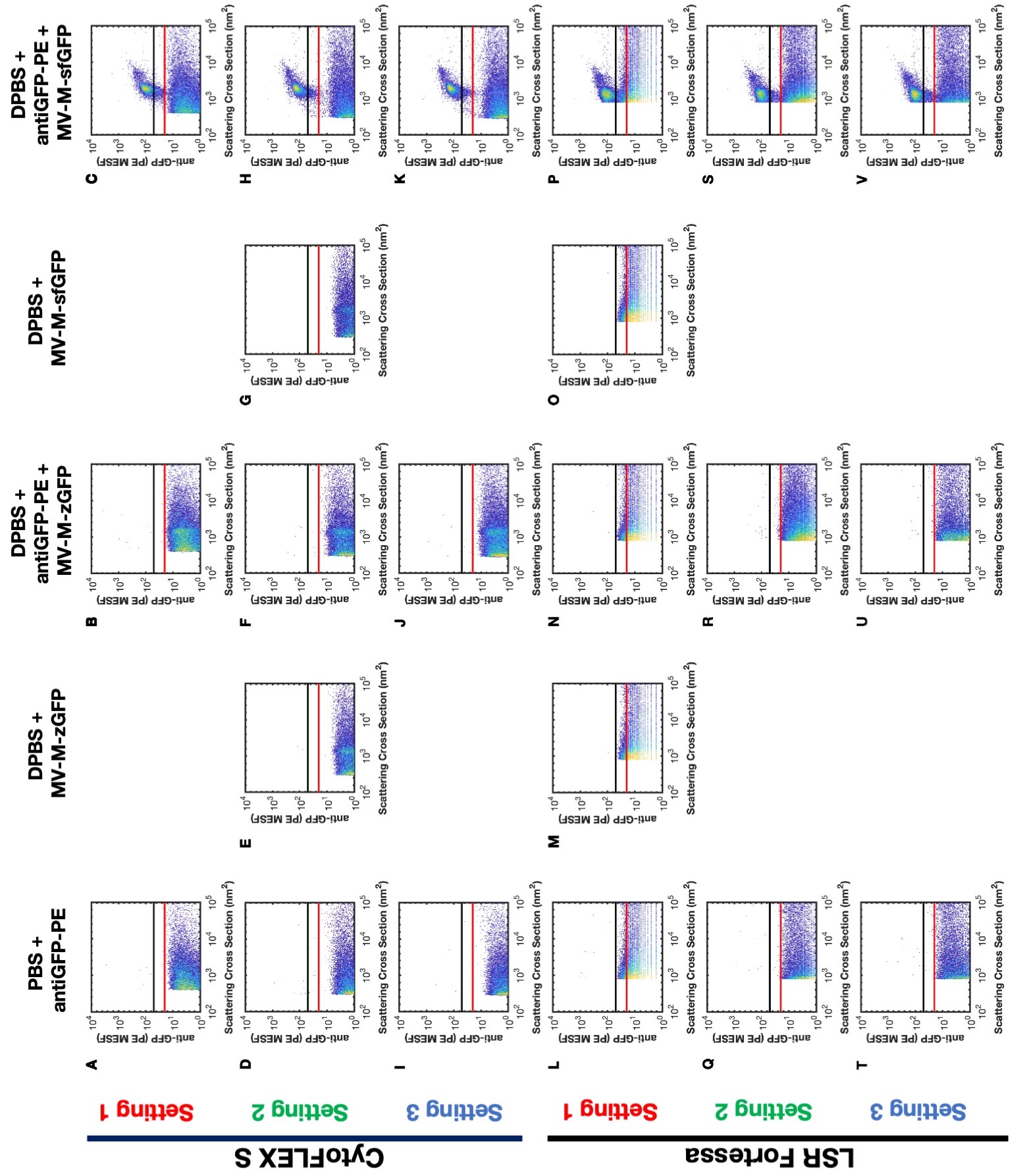
